# Mist4Cores: Reliable batch image stitching for dendrochronological cores

**DOI:** 10.1101/2025.10.17.667188

**Authors:** M García-Hidalgo, Y Ye, A García-Pedrero, JM Olano

**Affiliations:** Instituto Universitario de Gestión Forestal Sostenible (iuFOR), Escuela Universitaria de Ingeniería de la Industria Forestal, Agronómica y de la Bioenergía (EiFAB), Universidad de Valladolid, Soria, Spain; Department of Computer Architecture and Technology, Universidad Politécnica de Madrid, Madrid, Spain; Center for Biomedical Technology, Universidad Politécnica de Madrid, Madrid, Spain

**Keywords:** digitization, open-source, stitching

## Abstract

MIST4Cores automates the end-to-end process of image stitching for camera-based methods. This software allows reliable stitching with customizable user configuration and directory monitoring through overlap analysis and plugin-driven assembly. Each function in the software serves a distinct role in facilitating robust, reproducible, and user-friendly image stitching for scientific workflows. The combination of free open software tools for image acquisition, image stitching, and image analysis creates an open-source ecosystem to promote further research in the dendrochronological area.

## Introduction

Current developments on sample digitization are pushing dendrochronology into the digital imaging world. Flatbed scanners are widespread at many dendrochronological laboratories as affordable and easy-to-use systems for sample digitization. However, the limited resolution and pixel quality of flatbed scanners still maintains high dependency on the direct supervision of the physical sample during the dendrochronological analysis. Recent methodologies for digitization with camera-based systems reach great levels of resolution and pixel quality, effectively incorporating dendrochronology into the digital world (García-Hidalgo *et al*. 2022, Griffin *et al* 2021, WSL 2023).

Image capturing with digital cameras also entails certain challenges. In the digitization process through flatbed scanners, the photo-sensible sensor collects reflected light by the sample surface, and the scanning software generates a unique image for the whole sample. On the other hand, camera-based systems capture one image per shoot, requiring a consequent stitching process to generate a single image for the whole sample. This stitching process is commonly run through generalist software to create the resulting panorama image.

General stitching software tools (e.g., PTGui (New House Int. Serv. Rotterdam, The Netherlands), Photoshop (Adobe Inc. San Jose, CA, USA) or Image Composite Editor (Microsoft research, Redmond, WA, USA) are primarily aimed at users who focus on collecting panoramas from landscapes, which are captured with changes in the angle between the camera and the object of interest across images. Furthermore, these stitching methods for general consumer use tend to adapt individual pixel characteristics (e.g., size, value or bit depth) for image alignment prioritizing the creation of engaging images over the strict fidelity to the original single images. Although there are some ways for lens distortion correction (Adobe Inc. 2025), this phenomenon of distancing of fidelity has increased due to the current incorporation of default smoothing or enhancement filters, in many cases beyond the users’ control. The use of this general stitching software, therefore, requires several control steps to reduce artifact occurrences in the image and to improve the reliability of the whole digitized image for scientific applications.

The generation of panorama images (*e*.*g*., Whole Slide Images) from individual shoots is also a recurrent problem in different scientific areas (Mohammadi *et al*. 2024). Opposite to the case of generalist stitching methods, reliability and replicability of results from the sample to the panorama output image are critical. In fact, open-source community has explored diverse reliable stitching solutions promoting user interaction and fidelity control about the entire process. These methods are usually focused on microscopic slides which maintain uniform focal distance and fixed displacement across single images.

Here we present MIST4Cores, an open-source software developed for image stitching of sequential macroscopic images from cores by camera-based systems. MIST4Cores automates image stitching through an easy-to-use graphical interface, and its batch processing mode substantially reduces time and effort for image generation. Furthermore, MIST4Cores ensures image reliability and traceability with a log file per each whole image output.

## Application description

MIST4Cores software automates the stitching of macroscopic images using a combination of methods in just a one-task tool. In a first step, OpenCV (Bradski & Kaehler 2000) automatically detects the overlapping percentage in consecutive images. Afterwards, MIST (Chalfoun *et al*. 2017) is responsible to stitch the pairs of consecutive images under a backing process in Fiji/ImageJ environment (Schindelin *et al*. 2012). In a third step, the complete stitched image is exported according to user requirements. MIST4Cores is written in Python (van Rossum & Drake 2009), an increasingly common programming language in the scientific field, which ensures readability and further development by potential users according to their needs. Finally, the software is also compiled as an executable for its use for only-users. The main steps in the pipeline and the role of each function are outlined below:

### 1. Computing Environment

In the first run of the software, MIST4Cores prepares system dependencies, Fig.1. A Configuration window must be filled according to user settings. User enters the ImageJ/Fiji location, the source folder of single images (“*Monitoring folder*”), which can be a hierarchical nested. MIST4Cores will stitch single images in groups according to the immediate parent folder from the image files. Thus, a robust pipeline for folder/sample managing facilitates the complete process. The user can also enter the desired output folder path and the output format. Available outputs are TIFF and OME.TIFF (Linkert *et al*. 2010), which are not limited by pixel number as JPG files are. The user can also include the maximum of RAM available and the estimated resolution in dots per inch of the final image. The Configuration settings are then saved to be automatically loaded in further runs by MIST4Cores. This file collects also the stitching settings for MIST (e.g. file pattern, input image path, output image path) which can be customized by the user, see Supplementary materials for a detailed description.

**Figure 1.**
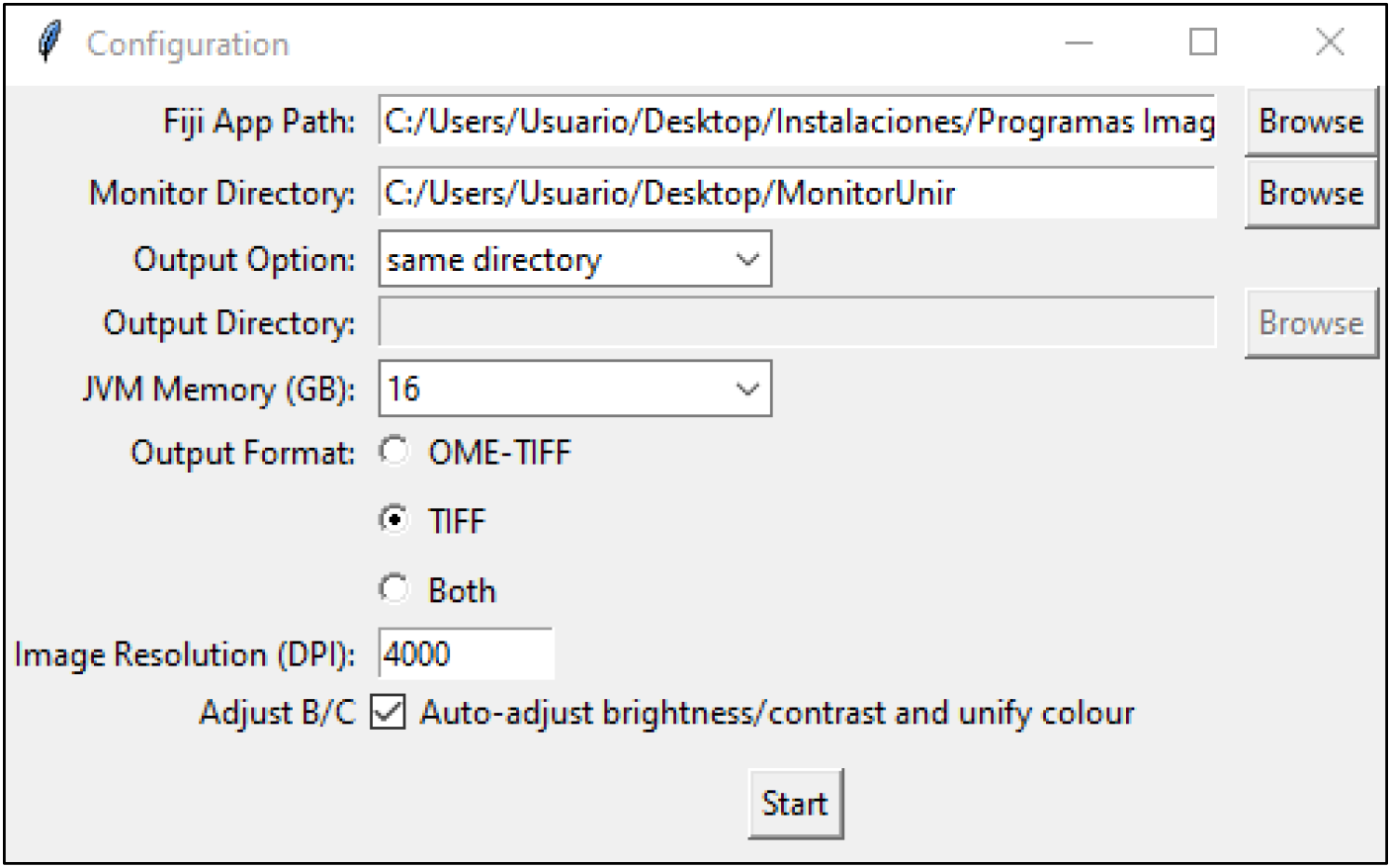
Configuration window of MIST4Cores.

### 2. Processing

MIST4Cores monitors user-specified directories for either pre-existing or newly created image folders which are then queued for processing. Within each folder, MIST4Cores identifies and sorts images matching the expected filename pattern, ensuring only files of interest are processed.

#### 2.1 Overlap Determination

Pairwise image overlaps are estimated using feature-based image registration (ORB keypoints and homography via OpenCV), providing the required parameters for subsequent stitching through MIST processing.

#### 2.2 Stitching Execution and Output

Fiji’s MIST stitching plugin with the calculated overlap and user-specified parameters runs the assembly of large composite images. Outputs are generated in user-selected formats (OME-TIFF, TIFF, or both) and can be adjusted for resolution (DPI).

#### 2.3 Logging and Reporting

Each run is documented with a summary log and metadata files, recording processing outcomes, image lists, overlap metrics, and run configurations for transparency and reproducibility.

All this pipeline is run in the backend, so the OpenCV or Fiji/ImageJ based methods are automatically managed and do not require any interaction by the user. The user is informed via terminal window to track the complete batching process in real time, in addition to the logging and reporting files generated at the end of each stitching process.

## Sample and Source Images

To maintain focus distance across the entire sample length, a totally flat surface is required for MIST4Cores adequate operation. A pre-processing step is thus critical to reduce variations of focal distance among single images. Once the tree core is mounted, a first precise sanding or horizontal cutting across the whole sample will ensure that both the basal mounting face and the surface of the core are completely parallel. The use of either a calibrating polishing machine or a cut with a sledge microtome is a must for a first core polishing. Successive finer grain polishing can be done with manual or mechanical alternatives, since it has little effect on surface depth.

MIST4Cores depends on established open-source tools for image processing (opencv) and scientific research (MIST), ensuring the fidelity among pixel values from sequential images for the stitching process. However, certain limitations appears in the longitudinal borders due to inherent methodological limitations of image padding (Zhang *et al*. 2023). In areas near the top or bottom edges of the image, the number of pixels in common between two consecutive images is reduced and, therefore, the stitching may suffer from slight blurring or displacement. Our analysis showed a displacement about 40±2 pixels (mean, SD) of discrepancies which is not generated in the central area, Figure 2. To reduce the concomitant padding and border artifacts in image stitching, we highly recommend to adapt the field of view in the shooting process. Samples of source images and the derived stitched complete images are provided in Zenodo.

**Figure 2.**
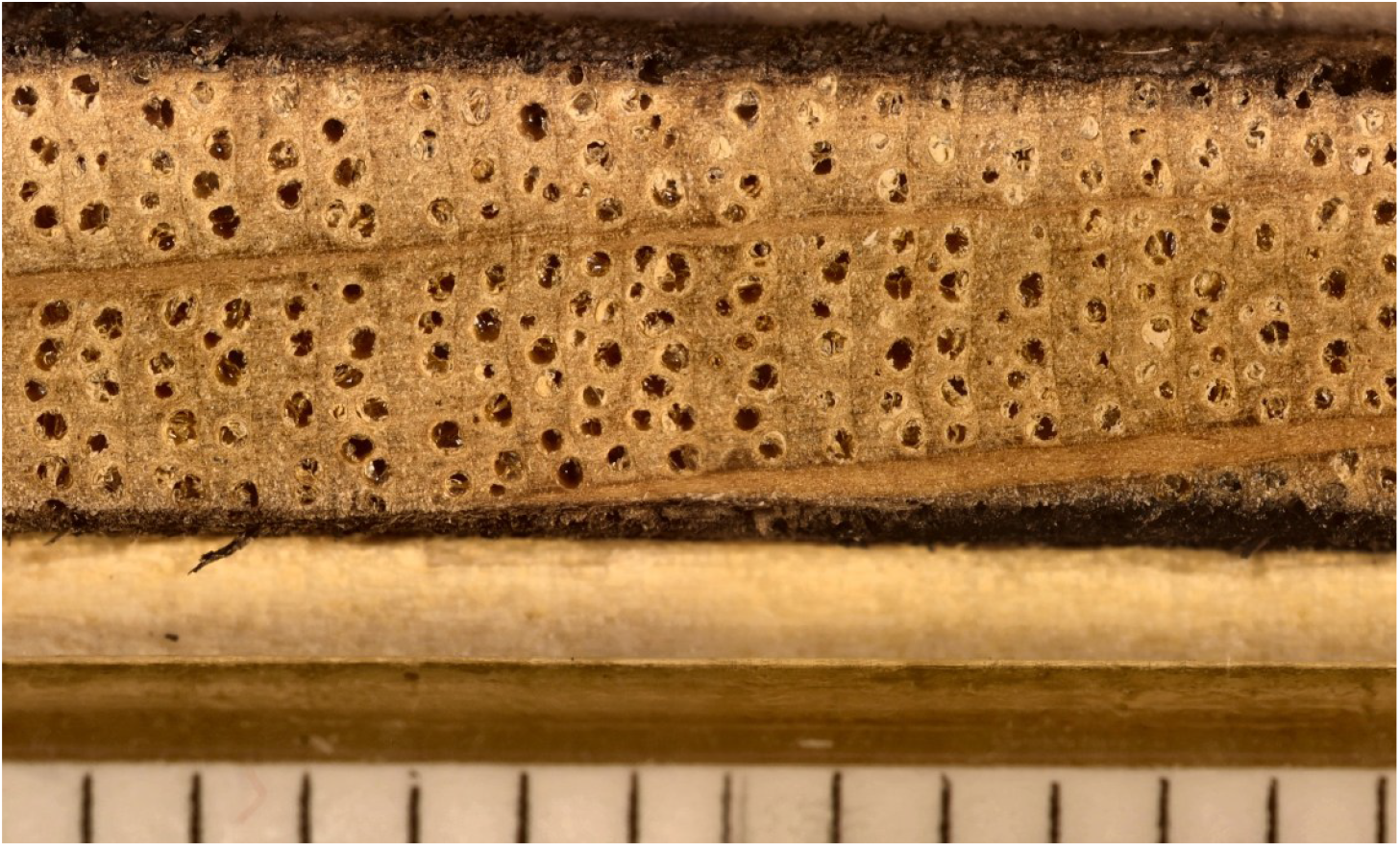
Resulting stitching detail. Bottom borders (physical scale) can show slight displacement in overlapping areas while the area of interest (core sample) maintains consistency.

## Computing Requirements

MIST4Cores is currently run using Java Virtual Machine (JVM) with limit assignation by the user. Since RAM availability limits computation reliability, we recommend to set half of available RAM for JVM. To ensure running stability, MIST4Cores would be run in systems with at least 32Gb of RAM without any other simultaneous processes running. Further developments for GPU computing could streamline stitching process while reducing costs.

## Code Availability

Application code/data and compiled release are available at https://github.com/cambiumrg and [Zenodo](https://doi.org/10.5281/zenodo.16831697).

