## Supplementary materials for "Mist4Cores: Reliable batch image stitching for dendrochronological cores"

### MIST4Cores

User Manual

V 1.0

Cambium Research Group 2025

A Python app for automating the stitching of images.

#### Installation

In the **Releases** section of MIST4Cores repository (https://github.com/cambiumrg) and Zenodo, you will find the executable file 'mist-stitcher.exe'. Once downloaded, open the program and follow the next steps of the user manual.

**Recommended:** Place the downloaded file in a new folder.

**Note:** You must have **Fiji** (Schindelin *et al.* 2012) installed beforehand (https://imagej.net/software/fiji/downloads), and also the plugin **MIST** (Chalfoun *et al.* 2017).

#### Initial Setup

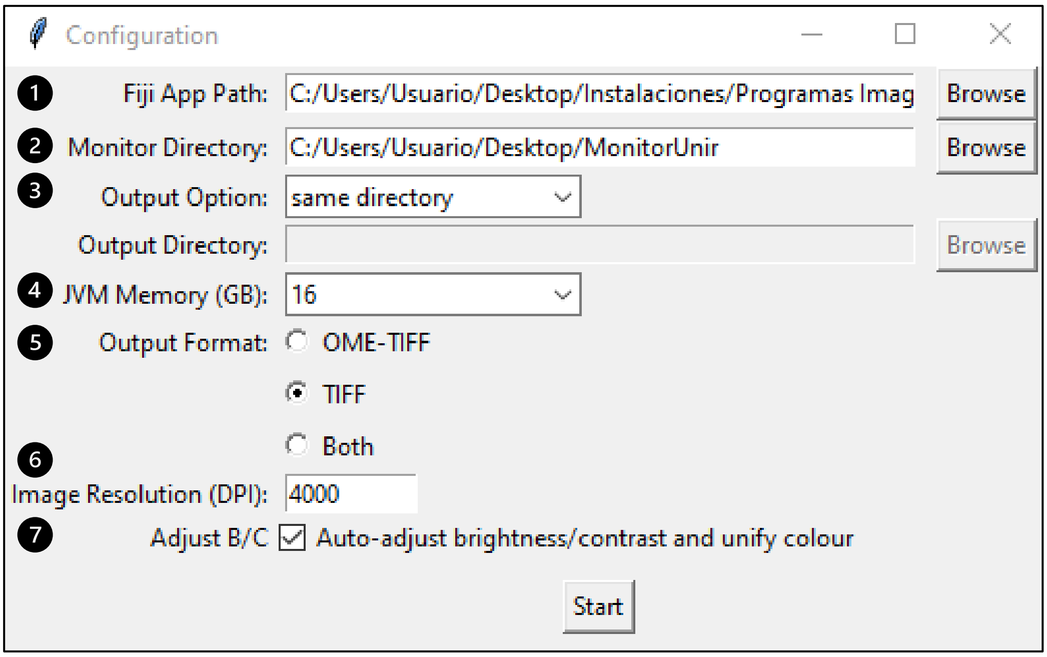


**Figure S1.** Configuration window. Each field is explained according to the reference numbers in for the initial set-up of MIST4Cores.

#### 1. Locate Fiji Installation

Specify the directory where Fiji Software (‘Fiji.app’) is located. E.g.: ‘C/Users/MainUser/Desktop/Fiji.app’ which includes Fiji executable file and required folders.

#### 2. Set the Monitoring Directory

Select the directory to be monitored for existing or new folders containing images. This directory may include a set of subfolders, each holding its own collection of images.

#### 3. Select Output Directory

- Choose ‘same directory’ to save the stitched image in the same directory as the original images.
- Choose ‘custom directory’ to select a custom output directory where all stitched images will be saved.

#### 4. Choose JVM memory size

- By default, the JVM memory size is set to 8 GB, but this value can be adjusted based on your system's available resources.
- If your machine has 32 GB of RAM, it is often advisable to allocate a smaller portion of memory to the JVM, such as 16 GB.

#### 5. Output Format

- Select the desired format for the stitched image: ‘.ome.tif’, ‘.tif’. In case that the user selects both, one file will be generated in each format.

#### 6. Image Resolution (DPI)

- Introduce an estimated dpi per image according to some image controls. This value will be saved in the output file to ensure compatibility with measuring software (e.g. CooRecorder (Cybis Elektronik & Data AB, Saltsjöbaden, Sweden).

This step does not replace the *Set-scale* or similar options of the measuring software.

**Note:** The DPI value is added to the .tif output metadata.

#### 7. Adjust Brightness / Contrast (Optional)

- When active, this mode reduces discrepancies on brightness and contrast among the single image series to create a seamless and visually coherent panorama.

#### Start Monitoring

After completing the configuration, you can choose between two options for stitching images:

- **'Process Existing Folders' (Recommended):** When the *'Save Mode'* button is pressed, the program will stitch all existing folders containing images, skipping those that have already been stitched, identified by the presence of files with "stitched" in their filename

- **'Monitor Future Folders':** The program will actively monitor new folders with images as they are added, ignoring any folders that already exist at the time this option is activated.

To stop monitoring and close the program, press the 'Stop' button.

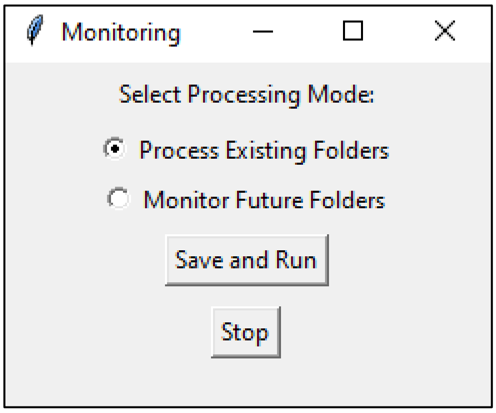


**Figure S2.** Monitoring Window. Stitching process can run on existing or future folders in the path set in the initial setup.

#### Images Extensions

MIST4Cores will process the monitoring folder for images with the following extensions: '.png', '.jpg', '.jpeg', and '.tif'.

#### Information File

Each stitched image has an associated file ('_info.txt') that contains details such as the sample name, stitching date, number of images stitched, horizontal and vertical overlap percentages, the input directory, the output directory, and the version of ImageJ used.

#### Log Summary

In the Monitor Directory you will find a file named `stitching_summary_log.txt` which contains the summary of the stitching process for all processed directories.

- Date and Time of Execution
- Directory
- Status:
    - "Successful stitching"
    - "Failed (Not all images selected)"

    - "Failed (Stitching image not saved)": usually because an Out of Memory exception is raised
    - "Failed"
- Time Elapsed (seconds)
- Images Processed
- Output Image Size (KB)

**Note:** Revise that the total number of images is correct. Not all images in a directory may be selected for stitching, even if the process completes successfully.

#### Configuration (Advanced)

Mist4Cores is totally customizable. One easy adjustment is to refine stitching parameters according to stitching methods or image name patterns.

##### MIST Parameters

The 'config.json' file contains the MIST stitching parameters, which can be modified directly within the file. This file is located in the same directory as the 'mist-stitcher.exe' executable.

Parameters accessed via 'config.get' file will use the saved configuration values. Other parameters are dynamically calculated at runtime.

```
{"fiji_app_path":"C:/Users/User/Desktop/Imaging/Fiji.app",
"monitor_directory": "C:/Users/Usuario/Desktop/MonitorUnir",
"output_option": "0",
"output_directory": null,
"memory_jvm_size_gb": "8",
"output_format": "tif",
"pattern": "([a-zA-Z0-9]+)_(\\d+)\\.(\\w+)",
"gridwidth": "14",
"gridheight": "1",
"starttile": "01",
"imagedir": "C:/Users/Usuario/Desktop/MonitorUnir/BA3CR02",
"filenamepattern": "BA3CR02_{pp}.jpg",
"filenamepatterntype": "SEQUENTIAL",
…
"loglevel": "MANDATORY",
"resolution_dpi": "6000",
"normalize_brightness": false
}
```

**Figure S3.** Part of config.json. The complete file contains MIST stitching parameters.

##### Image Name Pattern

The program uses the following regular expression to match image file names:

**([a-zA-Z0-9]+)_(\d+)\.(\w+)** which means that files have an alphanumeric prefix followed by an underscore '_' and two digits, ending with the file extension

Example: `GaQp01A15_01.jpg`

The underrun MIST method accepts the pattern defined as: prefix_{pp}.jpg

If you want to modify the pattern, you can do so by editing the 'config.json' file under the 'pattern' key.

**Note:** If the images do not match the defined pattern, the folder will be discarded, and the program will move on to the next one.
